# Museum genomics links *MC1R* alleles to adaptive winter coat color polymorphism in the long-tailed weasel

**DOI:** 10.64898/2025.12.17.694841

**Authors:** João Pimenta, Inês Miranda, Liliana Farelo, Marcela Alvarenga, L. Scott Mills, José Melo-Ferreira

## Abstract

Understanding the architecture of biological adaptations is a major endeavor of evolutionary biology. Using Natural History collections, we study the genetic basis and evolution of white/brown winter coat color variation in the long-tailed weasel (*Neogale frenata*), a crucial phenological adaptation for camouflage in habitats with seasonal snow. We produced whole-genome sequencing data for museum specimens, along two winter color morph transition areas in North America, at the West and East coasts. Genome-wide association scans identified a single genomic region linked to color variation polymorphism with approximately 300kb and 200kb in the West and East regions, respectively, which included the pigmentation gene *MC1R*. We identified three *MC1R* alleles, two of which with deletions of nine or eight amino acids, alternatively associated with the winter brown morphs in the West and East, respectively. These deletions affect the second transmembrane domain, and in one case also the first extracellular loop, which *in silico* analyses predicted to impact the protein’s function. Our findings show alternative intraspecific evolutionary solutions for environmental adaptation in long-tailed weasels, building on the evidence that major genes of the melanin production pathway are hotspots for recurrent and independent evolution of winter camouflage adaptation.

## Introduction

Seasonal coat color molt, from summer brown to winter white, is a remarkable phenological adaptation observed in more than 20 mammal and bird species, facilitating year-round crypsis in habitats subject to seasonal snow cover (Mills et al., 2018; Mills et al., 2013; Zimova et al., 2018). This phenological trait is primarily triggered by photoperiod (Zimova et al., 2018) and is predicted to be significantly impacted by climate change, given decreases in snow cover duration induced by global warming (Ferreira et al., 2023; Gottlieb & Mankin, 2024; Mills et al., 2013; Stokes et al., 2025). Winter white individuals may experience extended periods of color mismatch with the background, leading to increased predation and elevated mortality rates (Atmeh et al., 2018; Emmel et al., 2025; Imperio et al., 2013; Melin et al., 2020; Zimova et al., 2020; Zimova et al., 2016). Work in color polymorphic tawny owls (*Strix aluco*) has shown that camouflage is also advantageous for predators to avoid detection by prey (Perrault et al., 2023). Seasonal color-changing species also exhibit polymorphism in winter color (Mills et al., 2018). Within populations experiencing less or more ephemeral snow duration, individuals may retain a year-round brown coloration (Mills et al., 2018; Zimova et al., 2018). Translocation experiments (Rothschild & Lane, 1957; Vage et al., 2005; Zimova et al., 2018) and genomics work (Ferreira et al., 2023; Giska et al., 2019; Jones et al., 2018; Miranda et al., 2021; Tietgen et al., 2021) have shown that this winter coloration polymorphism is genetically determined. Given the threats to this adaptation posed by human-mediated climate change, studying the genetic basis and molecular mechanisms underlying this polymorphism is fundamental for understanding species’ potential for rapid adaptation (Ferreira et al., 2023; Mills et al., 2018; Zimova et al., 2018).

Studies in several species polymorphic for SCC have linked distinct loci to winter color variation. For example, in two hare species, snowshoe hares and mountain hares, it has been shown that the *Agouti Signaling Protein* (*ASIP*) underlies the trait polymorphism (Giska et al., 2019; Jones et al., 2018), while in white-tailed jackrabbits, seasonal color polymorphism is determined by variation at *Endothelin Receptor B* (*EDNRB*), *Corin Serine Peptidase* (*CORIN*), and *ASIP* genes (Ferreira et al., 2023). Alternatively, studies in the arctic fox and the least weasel have shown that the pigmentation gene *Melanocortin-1 Receptor* (*MC1R*) (Garcia-Borron et al., 2005) is associated with alternative winter coloration (Miranda et al., 2021; Vage et al., 2005). While in Arctic foxes two amino acid mutations underlie the polymorphism (Vage et al., 2005), in the least weasel a single amino acid change causes the winter brown coloration morph (Miranda et al., 2021; Miranda et al., 2025). Evidence for a selective sweep of winter brown genetic variants has been found for least weasels in northern Europe, possibly in response to snow cover retraction after the last glacial maximum (Miranda et al., 2021; Miranda et al., 2025); a similar sweep likely occurred in snowshoe hares (Jones et al., 2018; Jones et al., 2020a, 2020b).

Here, we dissect the genetic basis of the winter coat color polymorphism in the long-tailed weasel (*Neogale frenata*) (Fig. 1a). This species is widely distributed in North America, and its range extends to northern regions of South America (Mills et al., 2018). It belongs to genus *Neogale* (Patterson et al., 2021), sister to *Mustela*, in which the least weasel (*M. nivalis*) and the stoat (*M. erminea*) also undergo winter whitening. The distribution of winter morphs in long-tailed weasels follows a latitudinal cline and is correlated with winter snow duration (Mills et al., 2018), suggesting that the trait variation underlies adaptation to environmental gradients. We use whole-genome sequencing of Natural History Museum specimens across color morph transition zones to understand the determinants of winter coat color polymorphism in the species.

**Figure 1.**
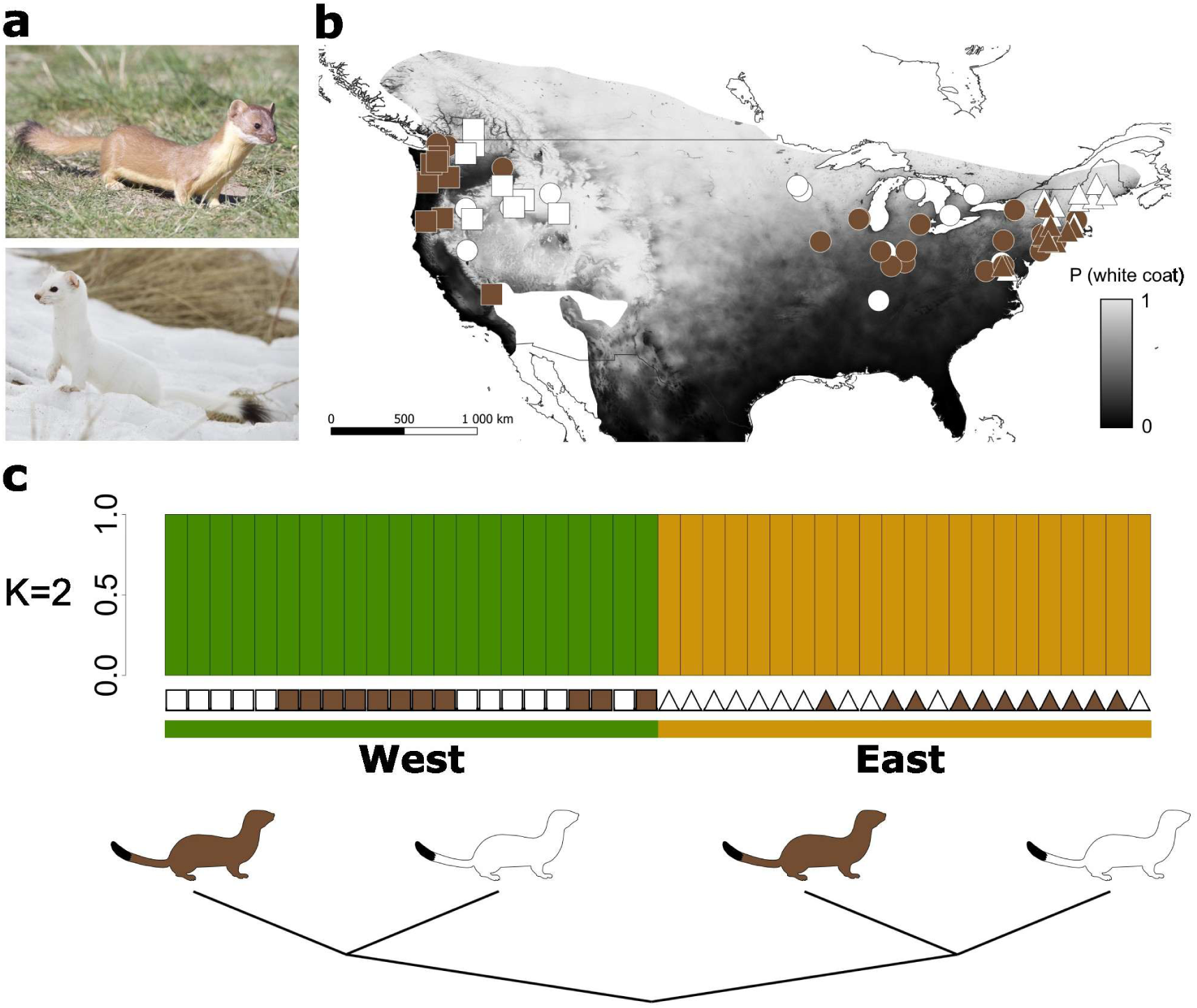
Coat color polymorphism and population structure in long-tailed weasel. (a) Coat color morphs of long-tailed weasel: brown (above) and white (below), John Krampl, iNaturalist, some rights reserved CC-BY. (b) The probability of the winter white color morph across the distribution of the species in North America, adapted from Mills et al. (2018). Data is accessible at https://doi.org/10.5061/dryad.8m0p1. Gradient regions represent the species distribution defined in the IUCN database (http://www.iucnredlist.org). Squares represent specimens included at least in the whole genome dataset for the west population. Triangles represent specimens included at least in the whole genome dataset for the east population. Circles represent samples included only in the genotyping dataset. Brown symbols represent winter brown samples, while white symbols represent winter white samples. (c) Above. Proportions of admixture inferred for K=2 ancestral clusters. For each region, samples are ordered based on their latitude (North to South) from left to right. Square colors represent winter phenotypes (brown – winter brown; white – winter white). Below. Maximum likelihood tree derived from allele frequencies estimated from unlinked genome-wide SNPs, with the American mink (*Neogale vison*) used as outgroup.

## Material and Methods

### Sampling and DNA extraction

A total of 102 long-tailed weasel (*Neogale frenata*) museum samples were included in this study, collected at the American Museum of Natural History (AMNH; New York), the Museum of Comparative Zoology (MCZ; Cambridge) and the Smithsonian National Museum of Natural History (USNM; Washington D.C.) in the USA (Table S1). These samples belong to two geographically distinct winter coat color transition regions in the USA and Canada, at the West (N=44) and East (N=58) coasts (Fig.1b and Table S1), and were originally collected between 1878 and 1964. All samples except one were collected during winter months, from November to March, and the winter phenotype for each sample was confirmed visually by examining the prepared skins. Notably, one of the samples was originally collected by Theodore Roosevelt, 26^th^ President of the USA in New York in 1874, and deposited in the collection of the National Museum of Natural History (accession nr. USNM 13507). Silica-based DNA extraction was performed on skin patches following Dabney et al. (2013).

### Library preparation and Whole-Genome Sequencing

Genomic DNA libraries were prepared for 44 samples included in analyses of whole-genome re-sequencing data (N=11 East brown, N=11 East White, N=11 West brown and N=11 West white). Double-indexed genomic libraries were prepared using a custom protocol derived from Meyer and Kircher (2010), including the USER (Uracil-Specific Excision Reagent) enzyme (New England BioLabs) in the blunt-end repair step (Kircher et al., 2012). Indexed libraries were quantified with quantitative polymerase chain reaction (qPCR). Finally, whole-genome low individual coverage data (targeting ∼1X per individual) was produced on an Illumina HiSeq X Ten (2 x 150 bp paired-end) at Novogene Co. Ltd. (Yuen Long, NT, Hong Kong).

### Raw Data Filtering

Raw sequencing data quality was assessed using FastQC, v.0.11.7 (Andrews, 2015), and adapter sequences were removed with Trimmomatic v.0.38 (Bolger et al., 2014) applying filters TRAILING:15, SLIDINGWINDOW:5:15, MINLEN:25. Overlapping pair-end reads were merged with PEAR v.0.9.1 (Zhang et al., 2014). Cleaned reads were mapped to the *Neogale vison* reference genome (Bioproject reference: PRJNA741394) (Karimi et al., 2022), using BWA-MEM v.0.7.17 (Li & Durbin, 2009) with default parameters and sorted with SAMtools v.1.13 (Li et al. 2009). PCR duplicates were removed by attributing distinct read groups to reads belonging to different flow cells using the function AddOrReplaceReadGroups and removing duplicated reads using the function MarkDuplicates from PicardTools v.2.18.17 (http://broadinstitute.github.io/picard/). Reads were locally realigned using RealignerTargetCreator and IndelRealigner from GATK v.3.8.1 (McKenna et al., 2010), and coverage statistics were retrieved with Qualimap v.2.2.1 (Okonechnikov et al., 2016). To avoid individual overrepresentation, samples with mean genome-wide coverage above 2X were down-sampled to a maximum of 1.5X using SAMtools v.1.13 (H. Li et al., 2009).

To reduce the potential error introduced by DNA damage present in museum historical samples (Briggs et al., 2007), in addition to the described use of the USER enzyme during library preparation, we applied an *in-silico* filter to remove C-to-T and G-to-A single-nucleotide polymorphisms (SNPs) (Axelsson et al., 2008). To apply the filter, we initially estimated allele frequencies on the entire whole-genome dataset (44 samples), using ANGSD v.0.940 (Korneliussen et al., 2014) with the following filters: -uniqueOnly, -GL 1, -remove_bads 1, -minQ 20 and -minMapQ 20. Then, sites with minor allele frequency 0.044 (i.e., 2 out of 88 chromosomes) for T or A were removed from the data set. To avoid representation biases, only sites represented by at least five individuals per coloration morph in both transition regions were considered for the following analyses.

### Population Structure

Population structure was assessed using principal component analysis (PCA) and a clustering assignment framework included in ANGSD (Korneliussen et al., 2014) and NGSadmix v.32 (Skotte et al., 2013), respectively, based on 123,718 unlinked genome-wide SNPs. The PCA was performed using the single-read approach implemented in ANGSD (-doIBS 2, -doCov 1), which is appropriate to low-coverage data. The SNP call was made using genotype likelihoods (-GL 1), with the following parameters: - minInd 22, -setMaxDepth 132, -doMaf 1, -minMaf 0.020 and -SNP_pval 1e-6. To minimize the effect of linkage disequilibrium, we used Plink v.1.90 (Purcell et al., 2007), applying the command –bp-space to keep SNPs distancing at least 20 kb. For each principal component, the eigenvectors and eigenvalues were estimated using the native function *eigen* from R v.4.3.1. Local population structure was also inspected for the Eastern and Western transition regions, based on 121,889 and 122,249 unlinked genome-wide SNPs respectively, using the same method described above, but adjusting to the lower sample sizes, using parameters -minInd 11, -setMaxDepth 66, -doMaf 1, and -minMaf 0.044. The cluster assignment analysis was performed with NGSadmix (Skotte et al 2013) using genotype likelihoods estimated with ANGSD (-GL 1, - doMajorMinor 1, -doMaf 1 and -doGlf 2), estimating admixture proportions for K=2 to K=4. A total of 200 independent runs were optimized using a maximum of 2000 Expectation-Maximization (EM) iterations, retaining the run with the best likelihood. In addition, a Neighbor-joining (NJ) tree was estimated using FastME v. 2.1.6.3 (Lefort et al., 2015) with the algorithm BioNJ, based on 123,718 unlinked genome-wide SNPs. Input files were generated from pairwise genetic distances estimated with ngsDist (Vieira et al., 2015), using genotype probabilities from ANGSD, with 100 bootstrap replicates and a block size of 25. Finally, the best-supported tree was obtained with RAxML (Stamatakis, 2014). To further examine the relationship between population structure and winter coloration morphs, we used TreeMix v.1.13 (Pickrell & Pritchard, 2012). Allele frequencies were determined for each morph-geography combination (East brown, East white, West brown, and West white) with ANGSD, using the American mink (*Neogale vison*) reference genome to determine the ancestral state (-GL 1, -doMaf 1, -doMajorMinor 5). Custom scripts were used to create the TreeMix input, and sites to include in the analysis were selected with a minimum distance of 20 kb between them, totaling 131,917 SNPs. The best tree topology was inferred by running 500 independent runs, with sample size correction disabled (-noss option), and a final global round of rearrangements (-global).

### Whole-genome scans of differentiation and association

Candidate regions for distinguishing winter color morphs were inferred using distinct approaches, taking advantage of the replication provided by the two morph transitions in the East and West regions. First, whole-genome scans of differentiation were performed with Popoolation2, v.1.201 (R. Kofler et al., 2011). Concatenated BAM files for each color morph were produced using SAMtools, v.1.6 (Heng Li et al., 2009), considering a minimum mapping and base quality of 20. The resulting mpileup file was then converted to sync format using Popoolation2, with a 5 bp window around inferred indels masked. We then performed a per-site Cochran–Mantel–Haenszel (CMH; McDonald (2009)) test based on 9,666,955 SNPs, identifying positions with shared allele frequency differences between color morphs in both transition regions, removing the confounding effect of putative population structure within each region. This analysis included filters --min-count 2 and --min-coverage 5. F_ST_ and Fisher’s Exact test (FET) estimations were also performed independently for each transition zone in non-overlapping windows of 20 kb, using the --karlsson-fst correction for F_ST_ and removing windows with a minimum covered fraction below 25% (--min-covered-fraction 0.25) and with less than 10 SNPs. Significance thresholds for CMH and FET were estimated using the Bonferroni correction for multiple tests (p-value < 0.05), and outlier regions of differentiation were defined based on the 99.9^th^ percentile of the F_ST_ values. The maximum coverage per pooled morph was set to three times the average coverage (--max-coverage 33).

An association case/control test was then performed with ANGSD using genotype likelihoods and testing for allele frequency differences between winter color morphs (-doAsso 1), pooling the same color morphs from both geographic regions. Allele frequencies were estimated from genotype likelihoods (-GL 1, -doMajorMinor 1 and - doMaf 1), for 7,363,078 SNPs, and sites were retained when represented in at least 50% of the individuals (-minInd 22), with a minimum of 50% and a maximum of three times the mean dataset coverage (-setMinDepth 22 -setMaxDepth 132). An association score-test (-doAsso 2) was performed using the eigenvectors of the first PC (PC1) of the local and global PCAs (described above) as a covariate, to account for the confounding effect of population structure (Skotte et al., 2012). The test was run using posterior probabilities (-doPost 1) estimated from genotype likelihoods (-GL1), following a dominant model, with the additional filters -minHigh 2 and -minCount 2. Results were summarized with a likelihood ratio statistic, converted to p-values by considering a chi-square distribution with one degree of freedom. Significance thresholds were estimated with a Bonferroni correction for multiple tests (p-value < 0.05).

Finally, we inspected the gene content of the region of elevated differentiation between winter color morphs and/or with significant association with the winter phenotype, which were consistently recovered across scans. We used the genome annotation for the *Neogale vison* reference genome to inspect the gene content of the association region. Additionally, associated regions were aligned against the annotated ferret (*Mustela putorius furo*) reference genome (Peng et al., 2014) using the function blastn from (Camacho et al., 2009), for additional inspection of candidate genes. We used the Ensembl annotation (https://www.ensembl.org/) and the gene predictions from the NCBI database (https://www.ncbi.nlm.nih.gov/) to characterize the gene content of the candidate-associated regions.

### SNP genotyping, MC1R variants, and local phylogeny estimation

To further validate the association regions identified in the genome scans, we genotyped selected SNPs in a larger sample dataset (N=85 specimens), belonging to the two independent transition regions and extending the sampling to the central region of the USA (Fig.1). For the candidate region in chromosome 7, we selected 54 SNPs showing high allele frequency differences between winter color morphs in each and/or in both transition regions. We independently estimated allele frequencies for each color morph and population using ANGSD, considering solely positions genotyped in at least five individuals (minInd 5) and with a maximum coverage of three times the population average (-setMaxDepth 33). Selected SNPs had allele frequency differences >0.5 between all pairs of winter color morphs. Genotyping was performed using the MassARRAY technology at the Genome Transcriptome Facility in the University of Bordeaux, France.

To facilitate visualization and statistical analysis, individuals with more than 30% of missing data were removed, followed by the removal of SNPs with more than 30% of missing data, resulting in a final dataset composed of 78 individuals and 36 SNPs. Finally, to detect positions with significant allele frequency changes between coat color morphs globally and within each transition region, we used the R package *SNPassoc* (González et al., 2007), with a Bonferroni-corrected significance value of p-value < 0.05.

Several previous works have reported a series of mutations occurring at the end of the second transmembrane domain and beginning of the first extracellular group of the *MC1R* protein, responsible not only for melanic phenotypes (McRobie et al., 2009) but also for winter coat color polymorphisms (Miranda et al., 2021; Vage et al., 2005). Therefore, a pair of primers was designed for this region to amplify a short fragment and evaluate if the same region of the gene could underlie the winter polymorphism in the long-tailed weasel (see Results). Three primers (10:1:10 ratio) were used in each PCR (Yang et al., 2015): a fluorescently labeled (6-FAM) M13 forward primer (5’-TGTAAAACGACGGCCAGT-3’), a sequence-specific forward primer with a 5’-M13 tail (5’-TGTAAAACGACGGCCAGTTGGCCGCGTCCGACCTG-3’), and a sequence-specific reverse primer (5’-ATGAGCACGTCGATGGCG-3’). Each reaction included 2.1µL of 2x QIAGEN Multiplex PCR Master Mix, 0.2µl of 10µM fluorescent forward primer, 0.2µL of 1µM forward primer, 0.2µL of 10µM reverse primer, 1.3 µl of water and 2µl of DNA. PCR conditions were 15 min denaturation at 95°C, 42 cycles of 30s at 95°C, 30s at 64°C, and 12s at 72°C, followed by 10min in final extension at 72°C. A size standard (GeneScan™ 500 LIZ®) was added to the fluorescent amplicons and fragments were separated by size on a capillary electrophoresis instrument (ABI 3500xl Genetic Analyzer).

The inferred *MC1R* variants, genotyped by PCR, were further analysed to explore their relationships by examining the genetic structure and phylogeny across this genomic region (25 kb upstream and downstream of the MC1R gene). A PCA was performed using the single-read approach implemented in ANGSD (-doIBS 2, -doCov 1), as described above. The SNP call was made using genotype likelihoods (-GL 1), with the following parameters: -minInd 11, -setMaxDepth 132, -doMaf 1, -minMaf 0.020 and - SNP_pval 1e-6. A NJ tree was also estimated following the same procedure described in the “Population Structure” section.

### Genetic diversity and selection

We estimated genome-wide nucleotide diversity (𝜋), Waterson’s theta (𝜃) and Tajima’s D with Popoolation, v.1.2.2 (Robert Kofler et al., 2011) for the four-coloration morph-geography groups. mpileup files were produced using the same procedure as for the whole-genome scans of differentiation and association. Diversity indexes were then estimated in windows of 50 kb and a step of 25 kb, considering a minimal covered fraction per window of 25%, a pool size of 22, a maximum coverage of three times the population average, and using the filters --min-count 2 and --min-coverage 5.

To test for signatures of selection along chromosome 7, we used a hidden Markov model approach as implemented in Pool-hmm v1.4.4 (Boitard et al., 2013), for pooled sequencing data (Boitard et al., 2012). For each morph-geography group, minimum coverage was set to 5, and the maximum coverage was set to 3 times the mean value (33X). The mean genome-wide theta (--theta) was calculated using Popoolation estimations. The transition probability between hidden states was set to values ranging between 1e-10 and 1e-30. The following settings were common to all analyses: -q 20 - e sanger.

### In silico inferences of the functional impact of MC1R mutations

The functional impact of the MC1R protein amino acid deletions found in winter brown morphs (see Results) was explored using *in silico* tools, SIFT (Hu & Ng, 2013) and CADD (Rentzsch et al., 2018). Due to the lack of available tools that predict the functional impact of insertion-deletions in non-model organisms, we used the human protein as a proxy to test the impact of the deletions. SIFT uses a heuristic method to find the most important features to predict the potential impact of 3n indels in gene function, applying a machine learning algorithm. CADD integrates diverse genomic annotations and uses machine learning methods to predict the functional consequences of genetic variants.

## Results

### Population structure follows the geographic distribution of the sampled specimens and not coloration morphs

Whole-genome sequencing of long-tailed weasels from the East and West coasts of the USA and Canada, representative of alternative winter morphs, resulted in an average sequencing coverage of 1.11X and 1.20X per specimen and 12.21X and 13.2X per winter morph for Western and Eastern geographic regions, respectively (Table S2). PCA and ancestral clustering showed that geography rather than winter color phenotype determined the genetic structure in our dataset (Fig.1c; Fig.S1), separating Eastern and Western populations. Results from the NJ tree inferred from pairwise genetic distances presented similar results, with geography as the main determinant for the genetic structure (Fig.S1b). Similarly, the population tree separated specimens from each region (Fig.1c). At the local geographical scale, PCAs showed shallow genetic structure across the Eastern and Western transition regions (Fig.S2).

### Winter coat color polymorphism is associated with a genomic region containing the MC1R gene

Taking advantage of the two replicated transition regions between color morphs, the Cochran-Mantel-Haenszel (CMH) association test identified a single genomic region significantly associated with the winter color morphs on chromosome 7 (Fig.2b). The result was consistent with the global case-control test based on genotype likelihoods (Fig.2a). The same genomic region showed significant differentiation when analyzing each transition region individually, using Fisher exact test (FET) and F_ST_ genome-wide scans (Fig.S3), which revealed a longer stretch of elevated differentiation in the Western (∼300kb) population than in the Eastern population (∼200kb). These results point to a genomic region of approximately 300 kb of elevated differentiation in chromosome 7 (scaffold coordinates 25,000 - 325,000 bp) as a candidate to underlie winter coat color polymorphism in the long-tailed weasel (Fig.S3f). The inspection of the reference genome annotation for the association region allowed identifying 11 genes: *PRDM7*, 5-hydroxyisourate hydrolase-like protein, *GAS8*, *DBNDD1*, *AFG3*-like protein, *DEF8*, tubulin beta-3 chain like protein, *TCF25*, *SPIRE2*, *FANCA*, and *ZNF276*. In the homologous region in the annotated ferret genome, which is located in scaffold GL896939.1 (44,000 - 329,000 bp, Peng et al. (2014)), 12 genes are included: *PRDM9* (instead of *PRDM7*), *URAH*, *GAS8*, *DBNDD1*, *AFG3L1P*, *DEF8*, *TUBB3*,

*MC1R*, *TCF25*, *SPIRE2*, *FANCA*, and *ZNF276*. Notably, the annotation of the *Neogale vison* reference genome does not include the *MC1R* gene sequence. Genotyping 36 SNPs along the association region in 78 specimens (Fig.S4, Table S3) showed that 22 and 15 SNPs were significantly differentiated (P < 0.05, Bonferroni-corrected) between color morphs in the Western and Eastern populations, respectively (Table S3). Of these, 7 SNPs showed significant allele frequency changes common to both transition regions (Table S3).

No signals of positive selection across the association region were inferred by Pool-HMM. Accordingly, we did not find reduced Tajima’s D or genetic diversity in the association region (Fig.S6). Nevertheless, we found slightly reduced genetic diversity in white individuals in comparison with brown individuals in the association region for the Western populations (Fig.S6).

### Polymorphic insertion-deletions in the coding region of the MC1R gene are associated with winter brown coloration

Sanger-sequencing of a region of *MC1R* that encompassed the second transmembrane domain and the first extracellular loop of the protein showed that winter brown specimens were homozygous or heterozygous for sets of amino acid deletions in the *MC1R* coding region, compared to the protein sequence of white winter specimens (Fig.3a-c, Fig.S5, and Fig.S7a). In Western winter brown samples, two deletions of 6 (T95_L100del) and 3 amino acids (A105_A107del or A106_A108del) were found in the second transmembrane domain and the first extracellular loop, respectively, considering the 2D structure of MC1R inferred from the human protein (Ringholm et al. 2004). In Eastern winter brown specimens, we found a single deletion of 8 amino acids (T95_E102del) in the second transmembrane domain (Fig.3b and c, and Fig.S7a). The amplified PCR fragments showed that winter brown samples were homozygous or heterozygous for the mutated allele, while winter white specimens were homozygous for the ancestral *MC1R* allele (Fig.S5b). Tests of functional impact suggested that both sets of deletions have a predicted impact on the protein function (Fig.S7b and c).

Local PCA analyses across the region of *MC1R* showed that PC1 (21.3% of variance), separates individuals by color morph, while PC2 (18.5% of variance) separates winter brown individuals from the East and West transition regions (Fig.3d). Congruently, the NJ tree (Fig.3e) showed a local phylogeny that clusters winter white individuals in a monophyletic group, placing the winter brown individuals in different branches of the tree.

## Discussion

This work took advantage of rich museum collections to investigate the genetic determination and evolution of winter color polymorphism in a species with seasonal camouflage. Analyzing sequencing data derived from low-quality and quantity DNA extracts from museum skins was challenging, yielding short sequence reads and increased duplication levels, resulting from the required PCR-based library preparations. This resulted in overall low individual whole genome sequencing coverages (1.11X). Yet, it also allowed recovering an average of 12.36X and 12.24X coverage per phenotype per transition zone, when considering sets of samples combining geography (east and west region, respectively) and winter coloration identified from the museum specimens (white or brown). By combining analyses based on genotype likelihoods, appropriate for low-coverage individual data, with those suitable for pools of individuals, the study controlled for individual representation, allowing for balanced pools.

The museum sampling covered two major areas of transition of winter coloration in long-tailed weasels, in the West and Eastern regions of the USA and Canada (Fig.1b). The distribution of winter coloration in the species corresponds to snow cover duration and ephemerality, suggesting that the coloration phenotypes are maintained by gradients of spatially varying selection related to camouflage against snow covered *versus* bare ground (Mills et al., 2018). Our population structure analyses of 123,718 unlinked SNPs revealed that genetic structure (Fig.1b and c) in our dataset was linked with geography rather than coloration morph. Such discordance between winter coloration and the demographic history of the populations has been shown in several other species, such as snowshoe hares (Jones et al., 2018), white-tailed jackrabbits (Ferreira et al., 2023), and least weasels (Miranda et al., 2021). This pattern suggests that adaptation along the environmental gradient is maintained by selection (Polechová & Barton, 2015). Even though our sampling represents the extremes of a presumably continuous distribution, which may have overestimated population structure, our results suggest that Eastern and Western populations are demographically independent replicates of the transition between winter color phenotypes, which provided a suitable context to use the transition zones to scan the genome for association with winter color (Miranda et al., 2021).

Our several genome scans, based on different criteria and datasets, such as differentiation and FET independently across each transition zone, or CMH and case-control association tests based on both transitions, suggest that a genomic region of 300 kb in chromosome 7 is associated with winter coloration in long-tailed weasels (Fig.2). In particular, the CMH test suggests that on the East and Western transitions, this genomic region shows synchronized allele frequency changes between white and brown winter morphs, at sites linked with the candidate causal variants. This was confirmed by the genotyping assay using an increased number of samples, which identified 7 SNPs with significant allele frequency changes in both transition zones, suggesting that the same genomic region underlies the phenotypic polymorphism across both transitions. This region contains *MC1R*, which is a well-known pigmentation gene involved in melanogenesis (Garcia-Borron et al., 2005) and has been widely shown to underlie color polymorphisms in nature (Hubbard et al., 2010), including seasonal coat color in the least weasels (Miranda et al., 2021). MC1R is a G-protein-coupled receptor that is predominantly expressed on the surface of melanocytes (Beaumont et al., 2005). When activated, typically by α-melanocyte-stimulating hormone (α-MSH), MC1R triggers eumelanin production, a dark brown pigment (Rouzaud et al., 2003). In contrast, in the presence of the agouti signaling protein (ASIP), the pigment production shifts to phaeomelanin, a yellow and red pigment, or pigment production is inhibited (Le Pape et al., 2009).

**Figure 2.**
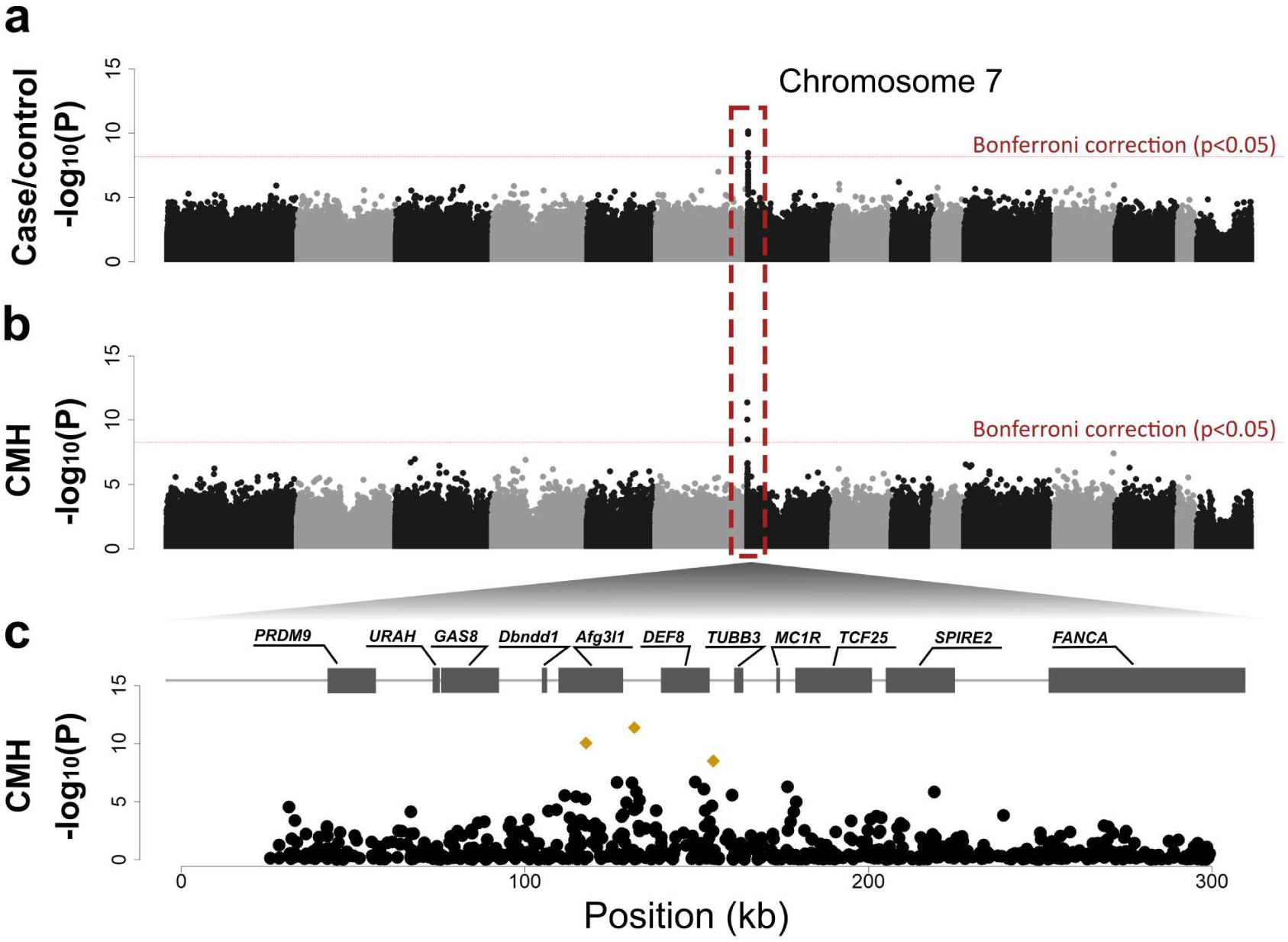
Genomic mapping of association for winter coloration morph in the long-tailed weasel. (a) Case–control association test of allele frequency differences between winter white and winter brown morphs for the complete dataset. The dashed red line indicates the Bonferroni-corrected threshold of 0.05. (b) CMH test of shared allele frequency differences between coloration morphs across the West and East populations. The dashed red line indicates the Bonferroni-corrected threshold of 0.05. (c) Zoom-in of the CMH test between coloration across the West and East populations. Genes annotated along the association region, based on the annotated ferret genome, are highlighted above the plot. Yellow polygons depict significantly associated SNPs.

**Figure 3.**
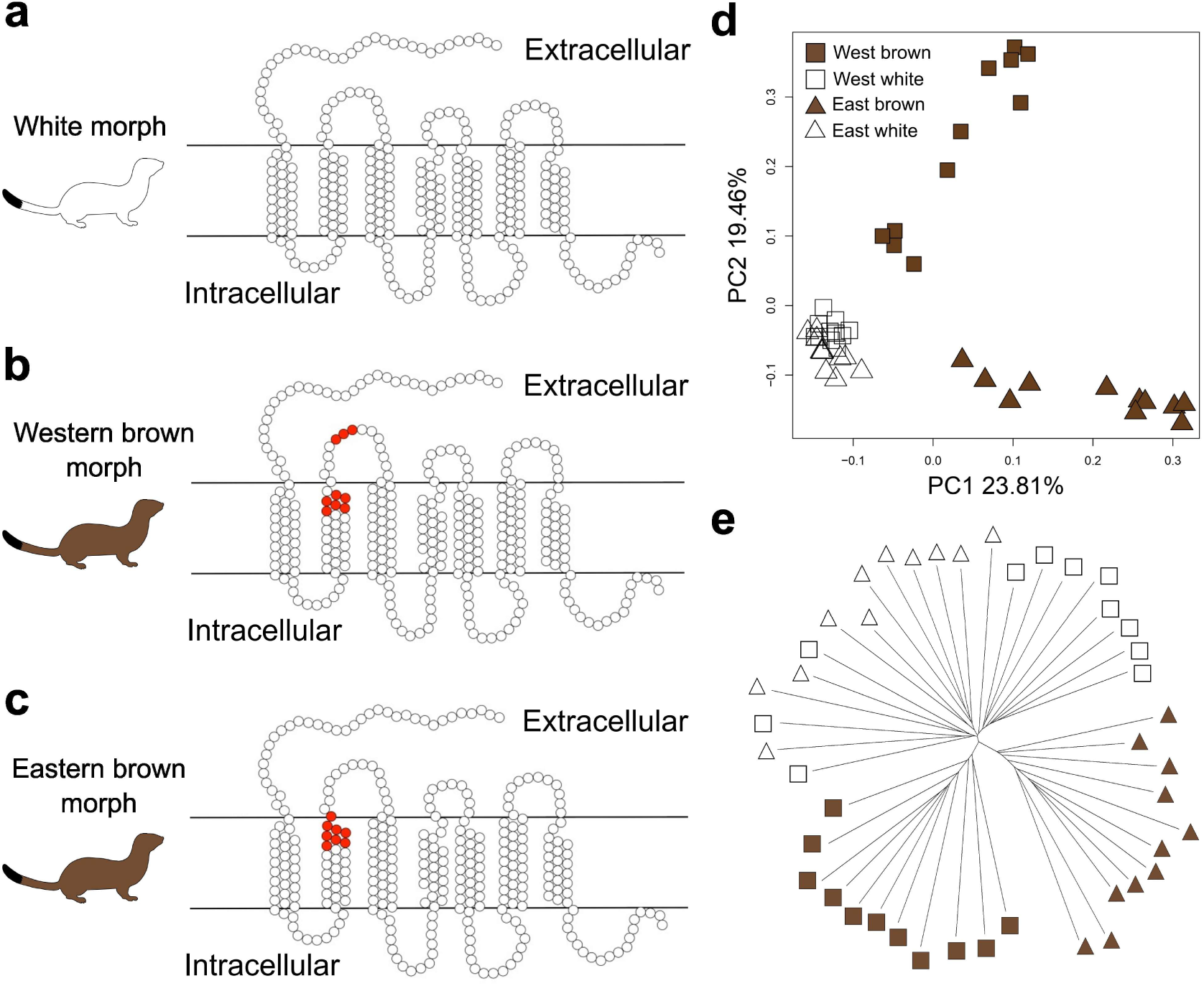
MC1R variants in the alternative winter coloration morphs in the long-tailed weasel. Two-dimensional (2D) representation of the MC1R variant associated with (a) the winter white coloration morph, (b) winter brown morph in the Western population, and (c) winter brown morph in the Eastern population. Red circles represent the deleted amino acids in the brown variants. (d) Principal component analysis of a 50kb region centred on *MC1R*, based on 220 SNPs. (e) Neighbor-joining tree of a 50kb region centered on the *MC1R* gene. Full symbols represent the winter brown morph and empty symbols the winter white phenotype. Squares correspond to individuals from the West population and triangles to the East population.

Our close inspection of the coding sequence of *MC1R*, using the PCR amplification of the target region (Fig.S5b), identified derived amino acid deletions perfectly associated with winter brown on both the East and West morph transitions. These results suggest a simple brown-dominant and white-recessive relationship of the alleles. Interestingly, different alleles were identified in each transition for the brown morphs, corresponding to different amino acid deletions, in an overlapping region of the protein, the second transmembrane domain and first extracellular loop (West), and the second transmembrane domain (East) (Fig.3b and c, and Fig.S5a). In the least weasel, Miranda et al. (2021) showed that an amino acid substitution at the transition between the second transmembrane domain and the first extracellular loop of *MC1R* is associated with the brown winter morph. Previously, a candidate gene approach in captive Arctic foxes and a whole-genome study in fallow deer showed that mutations causing amino acid changes in *MC1R* were associated with the blue and white morph phenotypes, respectively (Reiner et al., 2020; Vage et al., 2005). Also, McRobie et al. (2009) described a 24 bp deletion in the same gene region we found in our work that was associated with a melanic variant in the coat color of gray squirrels. Our *in silico* analyses suggested that the detected variants have an impact on the protein function (Fig.S7b and c). Overall, the segregation of the brown-associated alleles across the genotyped and phenotyped individuals suggests that winter coloration in long-tailed weasels has simple Mendelian inheritance with dominant brown, which agrees with findings in least weasels (Miranda et al., 2021).

While the functional consequence of the mutations identified in long-tailed weasels remains to be investigated, Miranda et al. (2025) showed that, in least weasels, an amino acid change in that region of MC1R interferes with the receptor’s ligand affinity to ASIP and αMSH. This change decreases MC1R response to both ligands, thus impacting the color switch during pelage molt, which can be used as a working hypothesis in this system as well. Particularly, mutated winter brown MC1R proteins may be less responsive to an expression peak of the *ASIP* gene known to occur during the autumn molt (Ferreira et al., 2017), which could render this expression peak insufficient to initiate the molt to white. Interestingly, in hares, *ASIP* has been implicated in adaptive camouflage evolution through cis-regulatory variation that alters the expression of the gene (Ferreira et al., 2023; Giska et al., 2019; Jones et al., 2018), suggesting that the *ASIP*-*MC1R* complex is a target of large effect for adaptive camouflage in mammals with seasonal coat color, but with specific routes for each species.

Our results show that different *MC1R* alleles are implicated in generating brown morphs in Eastern and Western regions of the distribution of long-tailed weasels. This questions whether the winter brown morph arose in the species once or multiple times independently. The existence of linked variation common to both morph transitions, with similar allele frequency changes from brown to white, suggests a single origin that may have diversified into the Eastern and Western alleles (Fig.2). However, further examining genetic variation within the associated region using local PCAs and NJ trees, revealed genetic similarity among winter white individuals but a clear distinction among winter brown morphs, particularly within the *MC1R* flanking regions (Fig.3d and e). Such increased variance and apparent basal position of the brown alleles may contradict the single-origin hypothesis, which may also reflect old origins maintained as balanced polymorphism in the species. In any case, our low coverage datasets provide little power to separate alternative hypotheses for the origin of the alleles, which should be investigated in future research.

Long-tailed weasels are both predators of small mammals and birds and prey for large carnivores, and the adaptive value of camouflage may thus come from both an increased ability to avoid predators and catch prey (Perrault et al., 2023). Given the predicted future decreases in snow cover extent caused by global warming (Brutel-Vuilmet et al., 2013), our findings have important implications for the understanding of the adaptive capacity of the species facing climate change. White morphs on habitats that are expected to lose winter snow cover in the future may face reduced fitness due to color mismatch (Ferreira et al., 2023; Mills et al., 2018; Perrault et al., 2023; Stokes et al., 2025). However, genetic polymorphism in winter coat color may promote evolutionary rescue to climate change, with the brown morph surviving better against a snowless background (Ferreira et al., 2023; Mills et al., 2018). Importantly, the dominant nature of the winter brown alleles in long-tailed weasels may have important implications for these future evolutionary dynamics. Ferreira et al. (2023) showed that the genetic architecture of winter brown influences the probability of future evolutionary rescue facing climate change, with recessive adaptive alleles (as found in snowshoe hares; Jones et al 2020a) leading to slower evolutionary responses when compared to additive brown alleles (as found in white-tailed jackrabbits; Ferreira et al. 2023). The dominant inheritance of the brown morph in long-tailed weasels could facilitate a faster response to selection, enhancing the potential for evolutionary rescue in populations exposed to shorter snow seasons.

While much of the existing genomic research on seasonal coloration has focused on hares (Ferreira et al., 2023; Giska et al., 2019; Giska et al., 2022; Jones et al., 2018; Jones et al., 2020a, 2020b; Mills et al., 2018), this study contributes to the growing understanding of the evolution of this adaptation in weasels and to identifying alternative evolutionary routes leading to camouflage adaptation across mammals. Future research should aim to further unravel the origins of the adaptive alleles and the species-specific dynamics of adaptation, providing deeper insights into the capacity of long-tailed weasels to adapt to the rapidly changing climate.

## Author Contributions

J.M.-F. coordinated the study; J.M.-F., L.S.M., J.P. designed research; J.M.-F. and L.SM. acquired funding; J.M.-F. sampled at Natural History collections; L.F. performed laboratory work; J.P., I.M., M.A. analyzed data; J.P., J.M.-F. wrote the paper; All authors contributed to the interpretation of the results, revised the paper and approved the final version.

## Acknowledgements

This work was supported by Fundação para a Ciência e a Tecnologia (FCT) (project grant to J.M.-F. under the ERC-Portugal programme). J.P. was supported by FCT Research Contract funded by project PTDC/BIA-EVL/28124/2017). I.M. was supported by FCT (research contract under the ERC-Portugal project). M.A. was supported by FCT (PhD Scholarship 2021.05642.BD and grant under the ERC-Portugal project). Further support was obtained from National Science Foundation grant DEB-1907022 to L.S.M. We thank the American Museum of Natural History, Museum of Natural Zoology, University of Harvard, and the National Museum of Natural History for granting sample loans, and Eleanor Hoeger, Breda Zimkus, Darrin Lunde, and Michael McGowen for support and sampling at museums. Additional support was obtained from the European Union’s Horizon 2020 Research and Innovation Programme under the Grant Agreement Number 857251.

## Bibliography

1. Andrews, S. (2015). FastQC: a quality control tool for high throughput sequence data. In

2. Atmeh, K., Andruszkiewicz, A., & Zub, K. (2018). Climate change is affecting mortality of weasels due to camouflage mismatch. Sci Rep, 8(1), 7648. 10.1038/s41598-018-26057-5

3. Axelsson, E., Willerslev, E., Gilbert, M. T. P., & Nielsen, R. (2008). The Effect of Ancient DNA Damage on Inferences of Demographic Histories. Molecular Biology and Evolution, 25(10), 2181–2187. 10.1093/molbev/msn163

4. Beaumont, K. A., Newton, R. A., Smit, D. J., Leonard, J. H., Stow, J. L., & Sturm, R. A. (2005). Altered cell surface expression of human MC1R variant receptor alleles associated with red hair and skin cancer risk. Human molecular genetics, 14(15), 2145–2154.

5. Boitard, S., Kofler, R., Françoise, P., Robelin, D., Schlötterer, C., & Futschik, A. (2013). Pool-hmm: a Python program for estimating the allele frequency spectrum and detecting selective sweeps from next generation sequencing of pooled samples. Mol Ecol Resour, 13(2), 337–340. 10.1111/1755-0998.12063

6. Boitard, S., Schlötterer, C., Nolte, V., Pandey, R. V., & Futschik, A. (2012). Detecting selective sweeps from pooled next-generation sequencing samples. Mol Biol Evol, 29(9), 2177–2186. 10.1093/molbev/mss090

7. Bolger, A. M., Lohse, M., & Usadel, B. (2014). Trimmomatic: a flexible trimmer for Illumina sequence data. Bioinformatics, 30(15), 2114–2120. 10.1093/bioinformatics/btu170

8. Briggs, A. W., Stenzel, U., Johnson, P. L., Green, R. E., Kelso, J., Prüfer, K., Meyer, M., Krause, J., Ronan, M. T., Lachmann, M., & Pääbo, S. (2007). Patterns of damage in genomic DNA sequences from a Neandertal. Proc Natl Acad Sci U S A, 104(37), 14616–14621. 10.1073/pnas.0704665104

9. Brutel-Vuilmet, C., Ménégoz, M., & Krinner, G. (2013). An analysis of present and future seasonal Northern Hemisphere land snow cover simulated by CMIP5 coupled climate models. The Cryosphere, 7(1), 67–80. 10.5194/tc-7-67-2013

10. Camacho, C., Coulouris, G., Avagyan, V., Ma, N., Papadopoulos, J., Bealer, K., & Madden, T. L. (2009). BLAST+: architecture and applications. BMC Bioinformatics, 10, 421. 10.1186/1471-2105-10-421

11. Dabney, J., Knapp, M., Glocke, I., Gansauge, M. T., Weihmann, A., Nickel, B., Valdiosera, C., Garcia, N., Paabo, S., Arsuaga, J. L., & Meyer, M. (2013). Complete mitochondrial genome sequence of a Middle Pleistocene cave bear reconstructed from ultrashort DNA fragments. Proc Natl Acad Sci U S A, 110(39), 15758–15763. 10.1073/pnas.1314445110

12. Emmel, A., Bickford, N., & Mills, L. S. (2025). Raptor Perception of Mismatch in Seasonally Polyphenic Prey. Am Nat, 206(6), 569–578. 10.1086/738016

13. Ferreira, M. S., Alves, P. C., Callahan, C. M., Marques, J. P., Mills, L. S., Good, J. M., & Melo-Ferreira, J. (2017). The transcriptional landscape of seasonal coat colour moult in the snowshoe hare. Mol Ecol, 26(16), 4173–4185. 10.1111/mec.14177

14. Ferreira, M. S., Thurman, T. J., Jones, M. R., Farelo, L., Kumar, A. V., Mortimer, S. M. E., Demboski, J. R., Mills, L. S., Alves, P. C., Melo-Ferreira, J., & Good, J. M. (2023). The evolution of white-tailed jackrabbit camouflage in response to past and future seasonal climates. Science, 379(6638), 1238–1242. 10.1126/science.ade3984

15. Garcia-Borron, J. C., Sanchez-Laorden, B. L., & Jimenez-Cervantes, C. (2005). Melanocortin-1 receptor structure and functional regulation. Pigment Cell Res, 18(6), 393–410. 10.1111/j.1600-0749.2005.00278.x

16. Giska, I., Farelo, L., Pimenta, J., Seixas, F. A., Ferreira, M. S., Marques, J. P., Miranda, I., Letty, J., Jenny, H., Hacklander, K., Magnussen, E., & Melo-Ferreira, J. (2019). Introgression drives repeated evolution of winter coat color polymorphism in hares. Proc Natl Acad Sci U S A, 116(48), 24150–24156. 10.1073/pnas.1910471116

17. Giska, I., Pimenta, J., Farelo, L., Boursot, P., Hacklander, K., Jenny, H., Reid, N., Montgomery, W. I., Prodohl, P. A., Alves, P. C., & Melo-Ferreira, J. (2022). The evolutionary pathways for local adaptation in mountain hares. Mol Ecol, 31(5), 1487–1503. 10.1111/mec.16338

18. González, J. R., Armengol, L., Solé, X., Guinó, E., Mercader, J. M., Estivill, X., & Moreno, V. (2007). SNPassoc: an R package to perform whole genome association studies. Bioinformatics, 23(5), 654–655. 10.1093/bioinformatics/btm025

19. Gottlieb, A. R., & Mankin, J. S. (2024). Evidence of human influence on Northern Hemisphere snow loss. Nature, 625(7994), 293–300. 10.1038/s41586-023-06794-y

20. Hu, J., & Ng, P. C. (2013). SIFT Indel: predictions for the functional effects of amino acid insertions/deletions in proteins. PLoS One, 8(10), e77940. 10.1371/journal.pone.0077940

21. Hubbard, J. K., Uy, J. A. C., Hauber, M. E., Hoekstra, H. E., & Safran, R. J. (2010). Vertebrate pigmentation: from underlying genes to adaptive function. Trends in Genetics, 26(5), 231–239.

22. Imperio, S., Bionda, R., Viterbi, R., & Provenzale, A. (2013). Climate change and human disturbance can lead to local extinction of Alpine rock ptarmigan: new insight from the western Italian Alps. PLoS One, 8(11), e81598. 10.1371/journal.pone.0081598

23. Jones, M. R., Mills, L. S., Alves, P. C., Callahan, C. M., Alves, J. M., Lafferty, D. J. R., Jiggins, F. M., Jensen, J. D., Melo-Ferreira, J., & Good, J. M. (2018). Adaptive introgression underlies polymorphic seasonal camouflage in snowshoe hares. Science, 360(6395), 1355–1358. 10.1126/science.aar5273

24. Jones, M. R., Mills, L. S., Jensen, J. D., & Good, J. M. (2020a). Convergent evolution of seasonal camouflage in response to reduced snow cover across the snowshoe hare range. Evolution, 74(9), 2033–2045. 10.1111/evo.13976

25. Jones, M. R., Mills, L. S., Jensen, J. D., & Good, J. M. (2020b). The Origin and Spread of Locally Adaptive Seasonal Camouflage in Snowshoe Hares. Am Nat, 196(3), 316–332. 10.1086/710022

26. Karimi, K., Do, D. N., Wang, J., Easley, J., Borzouie, S., Sargolzaei, M., Plastow, G., Wang, Z., & Miar, Y. (2022). A chromosome-level genome assembly reveals genomic characteristics of the American mink (Neogale vison). Commun Biol, 5(1), 1381. 10.1038/s42003-022-04341-5

27. Kircher, M., Sawyer, S., & Meyer, M. (2012). Double indexing overcomes inaccuracies in multiplex sequencing on the Illumina platform. Nucleic Acids Res, 40(1), e3. 10.1093/nar/gkr771

28. Kofler, R., Orozco-terWengel, P., De Maio, N., Pandey, R. V., Nolte, V., Futschik, A., Kosiol, C., & Schlötterer, C. (2011). PoPoolation: a toolbox for population genetic analysis of next generation sequencing data from pooled individuals. PLoS One, 6(1), e15925.

29. Kofler, R., Pandey, R. V., & Schlötterer, C. (2011). PoPoolation2: identifying differentiation between populations using sequencing of pooled DNA samples (Pool-Seq). Bioinformatics, 27(24), 3435–3436. 10.1093/bioinformatics/btr589

30. Korneliussen, T. S., Albrechtsen, A., & Nielsen, R. (2014). ANGSD: Analysis of Next Generation Sequencing Data. BMC Bioinformatics, 15(1), 356. 10.1186/s12859-014-0356-4

31. Le Pape, E., Passeron, T., Giubellino, A., Valencia, J. C., Wolber, R., & Hearing, V. J. (2009). Microarray analysis sheds light on the dedifferentiating role of agouti signal protein in murine melanocytes via the Mc1r. Proceedings of the National Academy of Sciences, 106(6), 1802–1807.

32. Lefort, V., Desper, R., & Gascuel, O. (2015). FastME 2.0: A Comprehensive, Accurate, and Fast Distance-Based Phylogeny Inference Program. Mol Biol Evol, 32(10), 2798–2800. 10.1093/molbev/msv150

33. Li, H., & Durbin, R. (2009). Fast and accurate short read alignment with Burrows-Wheeler transform. Bioinformatics, 25(14), 1754–1760. 10.1093/bioinformatics/btp324

34. Li, H., Handsaker, B., Wysoker, A., Fennell, T., Ruan, J., Homer, N., Marth, G., Abecasis, G., Durbin, R., & Genome Project Data Processing, S. (2009). The sequence alignment/map format and SAMtools. Bioinformatics, 25(16), 2078–2079. 10.1093/bioinformatics/btp352

35. Li, H., Handsaker, B., Wysoker, A., Fennell, T., Ruan, J., Homer, N., Marth, G., Abecasis, G., Durbin, R., & Subgroup, G. P. D. P. (2009). The sequence alignment/map format and SAMtools. Bioinformatics, 25(16), 2078–2079.

36. McDonald, J. H. (2009). Handbook of biological statistics (Vol. 2). sparky house publishing Baltimore, MD.

37. McKenna, A., Hanna, M., Banks, E., Sivachenko, A., Cibulskis, K., Kernytsky, A., Garimella, K., Altshuler, D., Gabriel, S., Daly, M., & DePristo, M. A. (2010). The Genome Analysis Toolkit: a MapReduce framework for analyzing next-generation DNA sequencing data. Genome Res, 20(9), 1297–1303. 10.1101/gr.107524.110

38. McRobie, H., Thomas, A., & Kelly, J. (2009). The genetic basis of melanism in the gray squirrel (Sciurus carolinensis). Journal of Heredity, 100(6), 709–714.

39. Melin, M., Mehtatalo, L., Helle, P., Ikonen, K., & Packalen, T. (2020). Decline of the boreal willow grouse (Lagopus lagopus) has been accelerated by more frequent snow-free springs. Sci Rep, 10(1), 6987. 10.1038/s41598-020-63993-7

40. Meyer, M., & Kircher, M. (2010). Illumina sequencing library preparation for highly multiplexed target capture and sequencing. Cold Spring Harb Protoc, 2010(6), pdb prot5448. 10.1101/pdb.prot5448

41. Mills, L. S., Bragina, E. V., Kumar, A. V., Zimova, M., Lafferty, D. J. R., Feltner, J., Davis, B. M., Hacklander, K., Alves, P. C., Good, J. M., Melo-Ferreira, J., Dietz, A., Abramov, A. V., Lopatina, N., & Fay, K. (2018). Winter color polymorphisms identify global hot spots for evolutionary rescue from climate change. Science, 359(6379), 1033–1036. 10.1126/science.aan8097

42. Mills, L. S., Zimova, M., Oyler, J., Running, S., Abatzoglou, J. T., & Lukacs, P. M. (2013). Camouflage mismatch in seasonal coat color due to decreased snow duration. Proc Natl Acad Sci U S A, 110(18), 7360–7365. 10.1073/pnas.1222724110

43. Miranda, I., Giska, I., Farelo, L., Pimenta, J., Zimova, M., Bryk, J., Dalen, L., Mills, L. S., Zub, K., & Melo-Ferreira, J. (2021). Museomics Dissects the Genetic Basis for Adaptive Seasonal Coloration in the Least Weasel. Mol Biol Evol, 38(10), 4388–4402. 10.1093/molbev/msab177

44. Miranda, I., Ruivo, R., Farelo, L., Alvarenga, M., Borowski, Z., Elmeros, M., Kalthoff, D. C., Merilä, J., Müller, J. P., Schley, L., Suchentrunk, F., Sundell, J., Rodrigues, M., Santos-Reis, M., Fernandes, C. R., Zub, K., Good, J. M., Mills, L. S., Castro, L. F. C.,…Melo-Ferreira, J. (2025). Ancient balanced polymorphism underlies long-standing adaptation for seasonal camouflage in the least weasel. bioRxiv, 2025.2011.2014.688436. 10.1101/2025.11.14.688436

45. Okonechnikov, K., Conesa, A., & Garcia-Alcalde, F. (2016). Qualimap 2: advanced multi-sample quality control for high-throughput sequencing data. Bioinformatics, 32(2), 292–294. 10.1093/bioinformatics/btv566

46. Patterson, B. D., Ramírez-Chaves, H. E., Vilela, J. F., Soares, A. E. R., & Grewe, F. (2021). On the nomenclature of the American clade of weasels (Carnivora: Mustelidae) [Review Article]. Journal of Animal Diversity, 3(2), 1–8. 10.52547/jad.2021.3.2.1

47. Peng, X., Alföldi, J., Gori, K., Eisfeld, A. J., Tyler, S. R., Tisoncik-Go, J., Brawand, D., Law, G. L., Skunca, N., & Hatta, M. (2014). The draft genome sequence of the ferret (Mustela putorius furo) facilitates study of human respiratory disease. Nature biotechnology, 32(12), 1250–1255.

48. Perrault, C., Morosinotto, C., Brommer, J. E., & Karell, P. (2023). Camouflage efficiency in a colour-polymorphic predator is dependent on environmental variation and snow presence in the wild. Ecology and Evolution, 13(12), e10824. 10.1002/ece3.10824

49. Pickrell, J., & Pritchard, J. (2012). Inference of population splits and mixtures from genome-wide allele frequency data. Nature Precedings, 1–1.

50. Polechová, J., & Barton, N. H. (2015). Limits to adaptation along environmental gradients. Proceedings of the National Academy of Sciences, 112(20), 6401–6406.

51. Purcell, S., Neale, B., Todd-Brown, K., Thomas, L., Ferreira, M. A., Bender, D., Maller, J., Sklar, P., de Bakker, P. I., Daly, M. J., & Sham, P. C. (2007). PLINK: a tool set for whole-genome association and population-based linkage analyses. Am J Hum Genet, 81(3), 559–575. 10.1086/519795

52. Reiner, G., Weber, T., Nietfeld, F., Fischer, D., Wurmser, C., Fries, R., & Willems, H. (2020). A genome-wide scan study identifies a single nucleotide substitution in MC1R gene associated with white coat colour in fallow deer (Dama dama). BMC genetics, 21, 1–8.

53. Rentzsch, P., Witten, D., Cooper, G. M., Shendure, J., & Kircher, M. (2018). CADD: predicting the deleteriousness of variants throughout the human genome. Nucleic Acids Research, 47(D1), D886–D894. 10.1093/nar/gky1016

54. Rothschild, M., & Lane, C. (1957). Note on change of pelage in the stoat (Mustela erminea L.). Proceedings of the Zoological Society of London,

55. Rouzaud, F., Annereau, J.-P., Valencia, J. C., Costin, G.-E., & Hearing, V. J. (2003). Regulation of melanocortin 1 receptor expression at the mRNA and protein levels by its natural agonist and antagonist. The FASEB journal, 17(14), 1–21.

56. Skotte, L., Korneliussen, T. S., & Albrechtsen, A. (2012). Association testing for next - generation sequencing data using score statistics. Genetic epidemiology, 36(5), 430–437.

57. Skotte, L., Korneliussen, T. S., & Albrechtsen, A. (2013). Estimating individual admixture proportions from next generation sequencing data. Genetics, 195(3), 693–702. 10.1534/genetics.113.154138

58. Stamatakis, A. (2014). RAxML version 8: a tool for phylogenetic analysis and post-analysis of large phylogenies. Bioinformatics, 30(9), 1312–1313. 10.1093/bioinformatics/btu033

59. Stokes, A. W., Hofmeester, T. R., Thorsen, N. H., Odden, J., Linnell, J. D. C., Zimova, M., & Pedersen, S. (2025). Mountain Hares Are Adapted to Historical Climates-Coat Colour Mismatch is Greatest in Areas With the Largest Reduction in Snow Cover Duration Over the Last 60 Years. Glob Chang Biol, 31(8), e70365. 10.1111/gcb.70365

60. Tietgen, L., Hagen, I. J., Kleven, O., Bernardi, C. D., Kvalnes, T., Noren, K., Hasselgren, M., Wallen, J. F., Angerbjorn, A., Landa, A., Eide, N. E., Flagstad, O., & Jensen, H. (2021). Fur colour in the Arctic fox: genetic architecture and consequences for fitness. Proc Biol Sci, 288(1959), 20211452. 10.1098/rspb.2021.1452

61. Vage, D. I., Fuglei, E., Snipstad, K., Beheim, J., Landsem, V. M., & Klungland, H. (2005). Two cysteine substitutions in the MC1R generate the blue variant of the Arctic fox (Alopex lagopus) and prevent expression of the white winter coat. Peptides, 26(10), 1814–1817. 10.1016/j.peptides.2004.11.040

62. Vieira, F. G., Lassalle, F., Korneliussen, T. S., & Fumagalli, M. (2015). Improving the estimation of genetic distances from Next-Generation Sequencing data. Biological Journal of the Linnean Society, 117(1), 139–149. 10.1111/bij.12511

63. Yang, Z., Steentoft, C., Hauge, C., Hansen, L., Thomsen, A. L., Niola, F., Vester-Christensen, M. B., Frödin, M., Clausen, H., Wandall, H. H., & Bennett, E. P. (2015). Fast and sensitive detection of indels induced by precise gene targeting. Nucleic Acids Res, 43(9), e59. 10.1093/nar/gkv126

64. Zhang, J., Kobert, K., Flouri, T., & Stamatakis, A. (2014). PEAR: a fast and accurate Illumina Paired-End reAd mergeR. Bioinformatics, 30(5), 614–620. 10.1093/bioinformatics/btt593

65. Zimova, M., Giery, S. T., Newey, S., Nowak, J. J., Spencer, M., & Mills, L. S. (2020). Lack of phenological shift leads to increased camouflage mismatch in mountain hares. Proc Biol Sci, 287(1941), 20201786. 10.1098/rspb.2020.1786

66. Zimova, M., Hacklander, K., Good, J. M., Melo-Ferreira, J., Alves, P. C., & Mills, L. S. (2018). Function and underlying mechanisms of seasonal colour moulting in mammals and birds: what keeps them changing in a warming world? Biol Rev Camb Philos Soc, 93(3), 1478–1498. 10.1111/brv.12405

67. Zimova, M., Mills, L. S., & Nowak, J. J. (2016). High fitness costs of climate change-induced camouflage mismatch. Ecol Lett, 19(3), 299–307. 10.1111/ele.12568

