## Supplementary Material for "Museum genomics links *MC1R* alleles to adaptive winter coat color polymorphism in the long-tailed weasel"

[1]CIBIO, Centro de Investigação em Biodiversidade e Recursos Genéticos, InBIO Laboratório Associado, Campus de Vairão, Universidade do Porto, 4485-661 Vairão, Portugal

[2]BIOPOLIS Program in Genomics, Biodiversity and Land Planning, CIBIO, Campus de Vairão, 4485-661 Vairão, Portugal

[3]Departamento de Biologia, Faculdade de Ciências, Universidade do Porto, 4169-007 Porto, Portugal

[4] Wildlife Biology Program, University of Montana, Missoula, MT 59812, USA

[5] Office of Research and Creative Scholarship, University of Montana, Missoula, MT 59812, USA

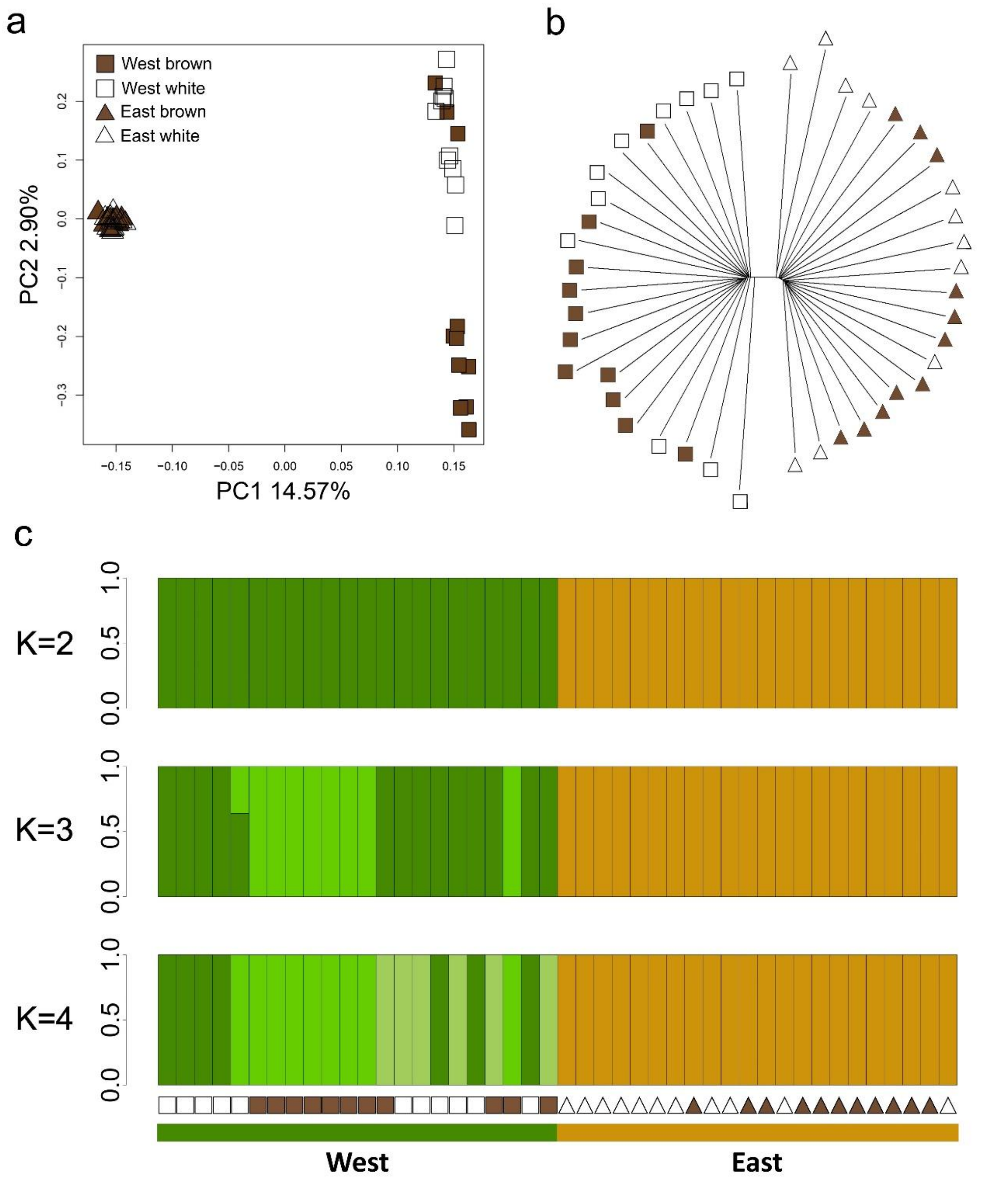


**Figure S1. Population structure in the long-tailed weasel.** (a) Principal component analysis of the complete whole-genome sequencing dataset of long-tailed weasels based on 115,002 SNPs. Brown polygons represent the brown morph, and white polygons represent the white morph. (b) Neighbour-joining tree of the complete dataset based on 114,102 SNPs. (c) Admixture proportions from 2 to 4 ancestral clusters (K). Squares represent Western specimens, while triangles represent Eastern samples. White symbols represent white winter morphs, while brown polygons represent brown winter morphs For each region, samples are ordered based on their latitude (North to South), from left to right.

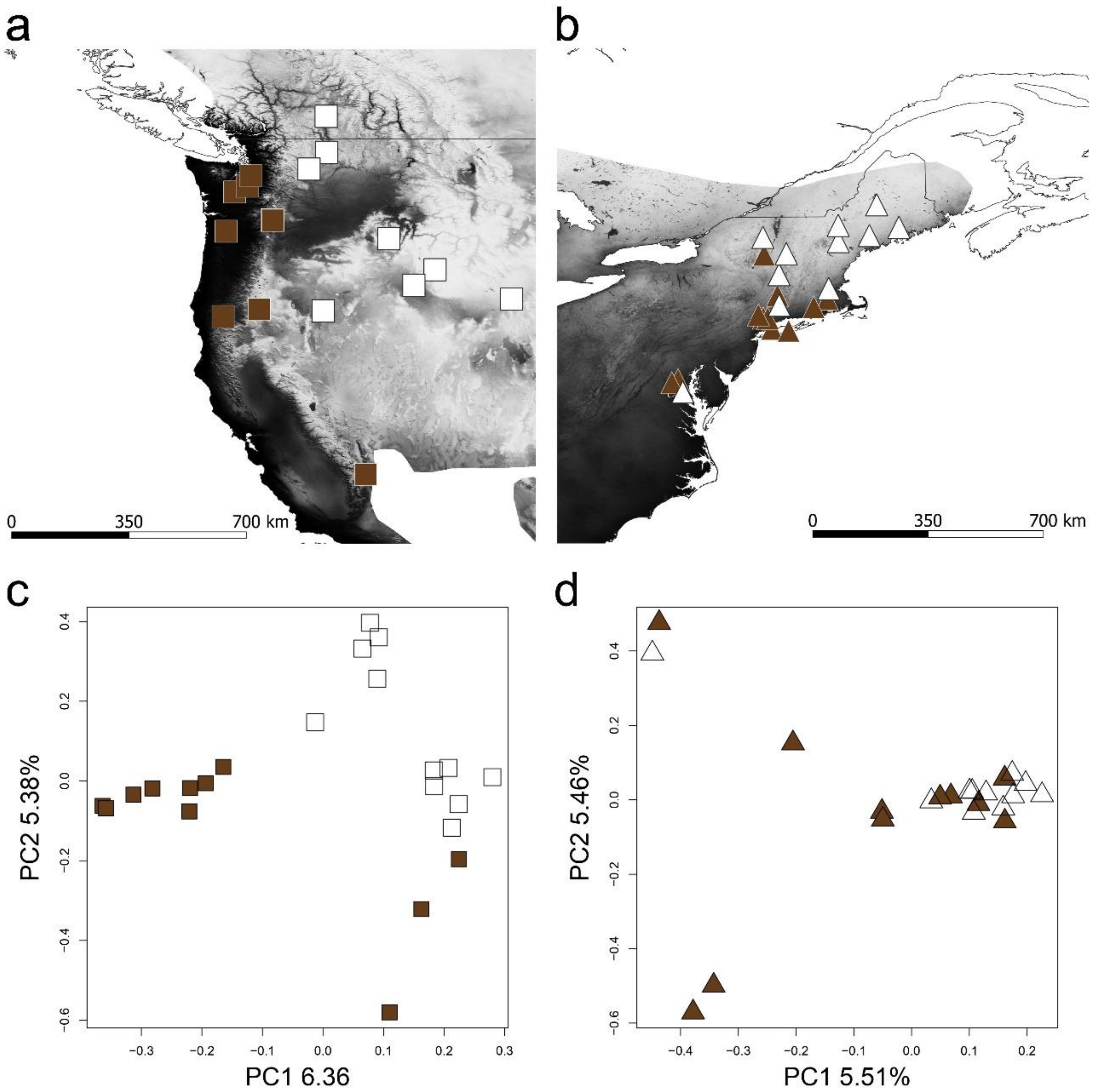


**Figure S2. Local population structure in Eastern and Western populations.** Geographic distribution of the whole genome dataset of long-tailed weasel across the (a) West and (b) East transition regions. Principal component analysis of genome-wide variation of (c) West and (d) East coast populations, based on 113,036 and 114,124 SNPs, respectively. Brown symbols represent the winter brown morph and white symbols represent the winter white morph.

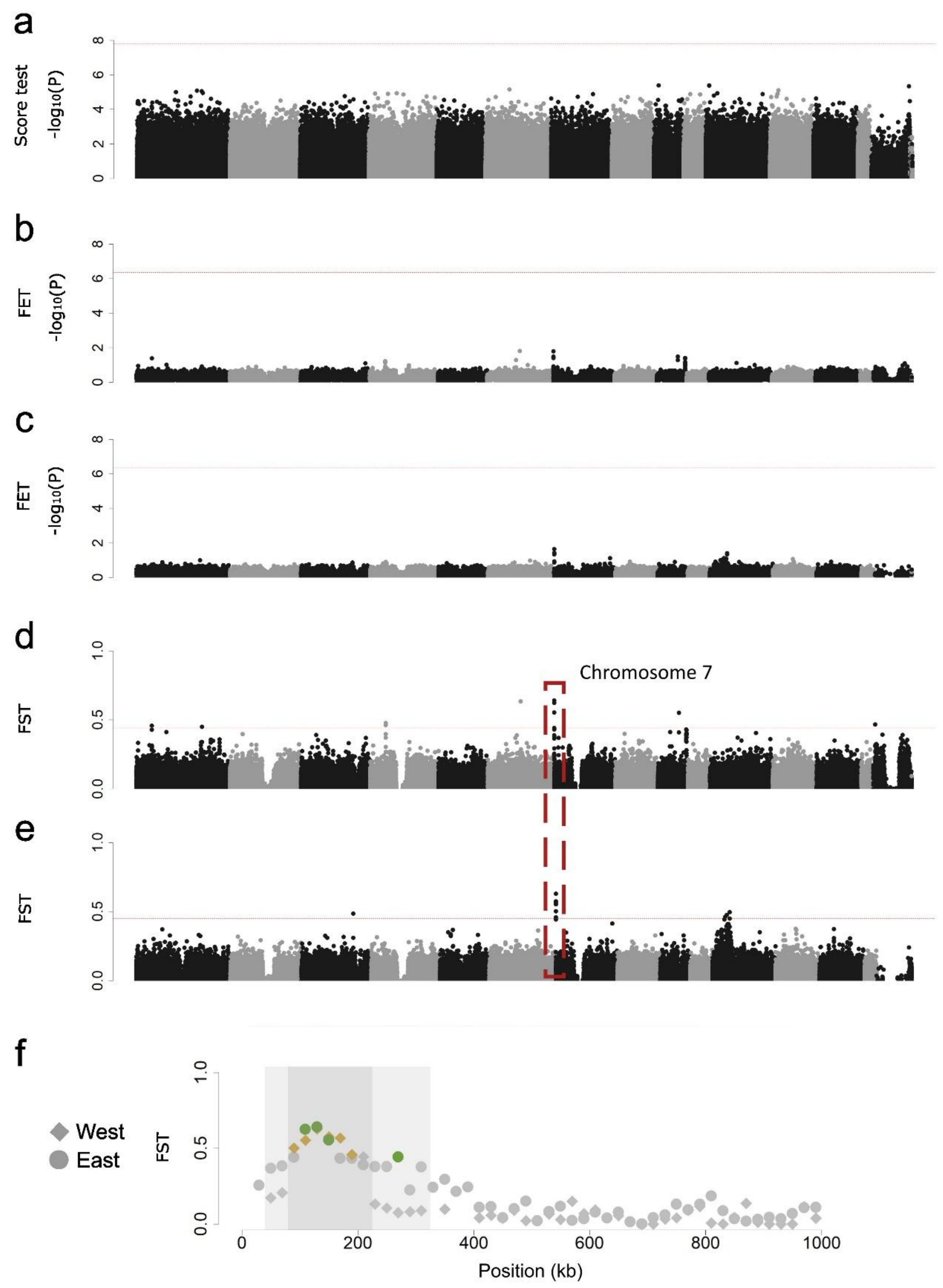


**Figure S3. Whole genome score test and local association scans.** (a) Score test of association between brown and white morphs for the entire dataset inferred from posterior genotype likelihoods. The test was controlled for population structure by including the first principal component of the global PCA (Figure S1a) as covariate. The dashed red line indicates the Bonferroni-corrected threshold of 0.05. (b-c) Fisher exact test (FET) of allele frequency differences between brown and white specimens for West (b) and East (c) populations, estimated from pooled data. Values are averaged in 20 kb non-overlapping windows. The dashed red line indicates the Bonferroni-corrected threshold of 0.05. (d-e) $F_{ST}$ divergence between brown and white winter colour morphs in West (d) and East (e) populations. Values are averaged in overlapping windows of 50 kb with a sliding window of 25 kb. The dashed red line represents the 99.9$^{th}$ percentile of windows with highest $F_{ST}$ values. (f) Zoom-in of $F_{ST}$ divergence estimation between coloration across the West and East populations. Colored polygons indicate windows above the 99.9$^{th}$ percentile of windows with highest $F_{ST}$ values. Grey shaded sections highlight the genomic region associated with the coat color polymorphism (light grey – west population – and dark grey – east population).

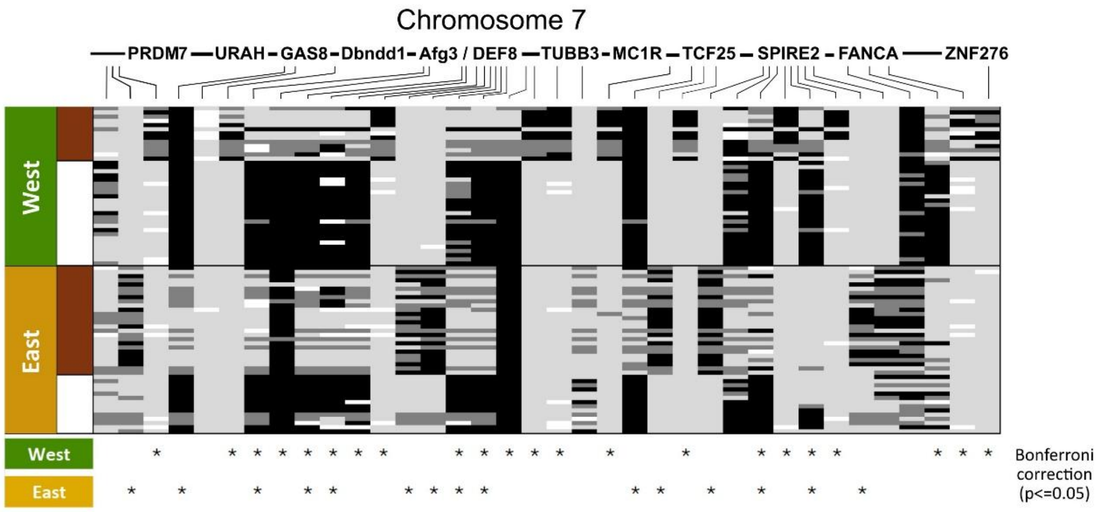


**Figure S4. SNP genotyping across the association region.** Genotypes for 36 SNPs along the candidate region (chromosome 7) in 78 individuals. Each row represents a specimen and each column a genotyped locus. The relative distribution of the SNPs along the association regions is shown. The left column indicates the population of origin (green for the West population and yellow for the East population) and coloration morphs (brown for the brown morph and white for the white morph). For each locus, and using *N. vison* genome as reference, genotypes are colored as black - homozygous derived allele; light grey - homozygous for the ancestral variant; dark grey - heterozygous; white - missing data. Below. The indication of positions with significant allele frequencies differences between winter morphs within both geographic regions estimated using a Bonferroni correction ($p < 0.05$).

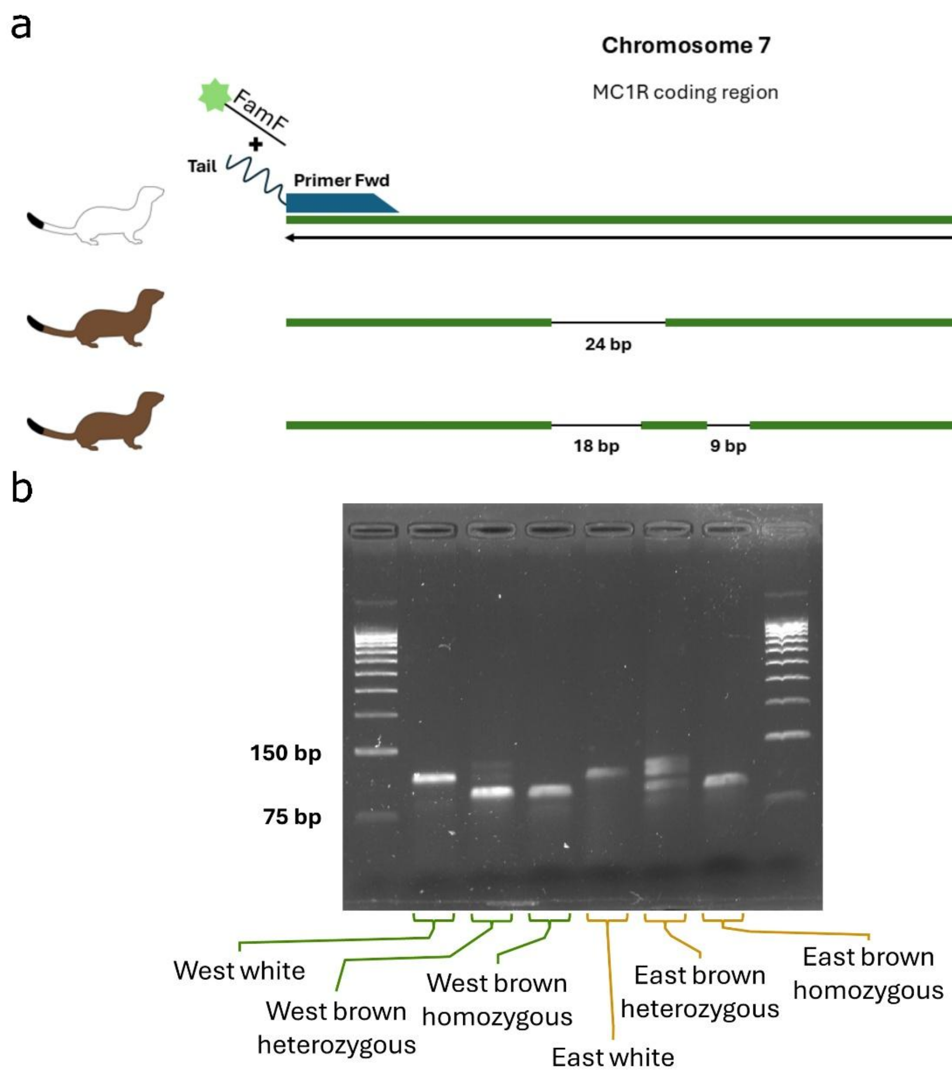


**Figure S5. Structural variants in *MC1R*.** (a) Scheme of the PCR genotyping strategy of the *MC1R* variants in the winter morphs for both transition regions using fluorescently labelled primers. (b) Agarose gel showing the genotype of the alternative variants of the *MC1R* gene.

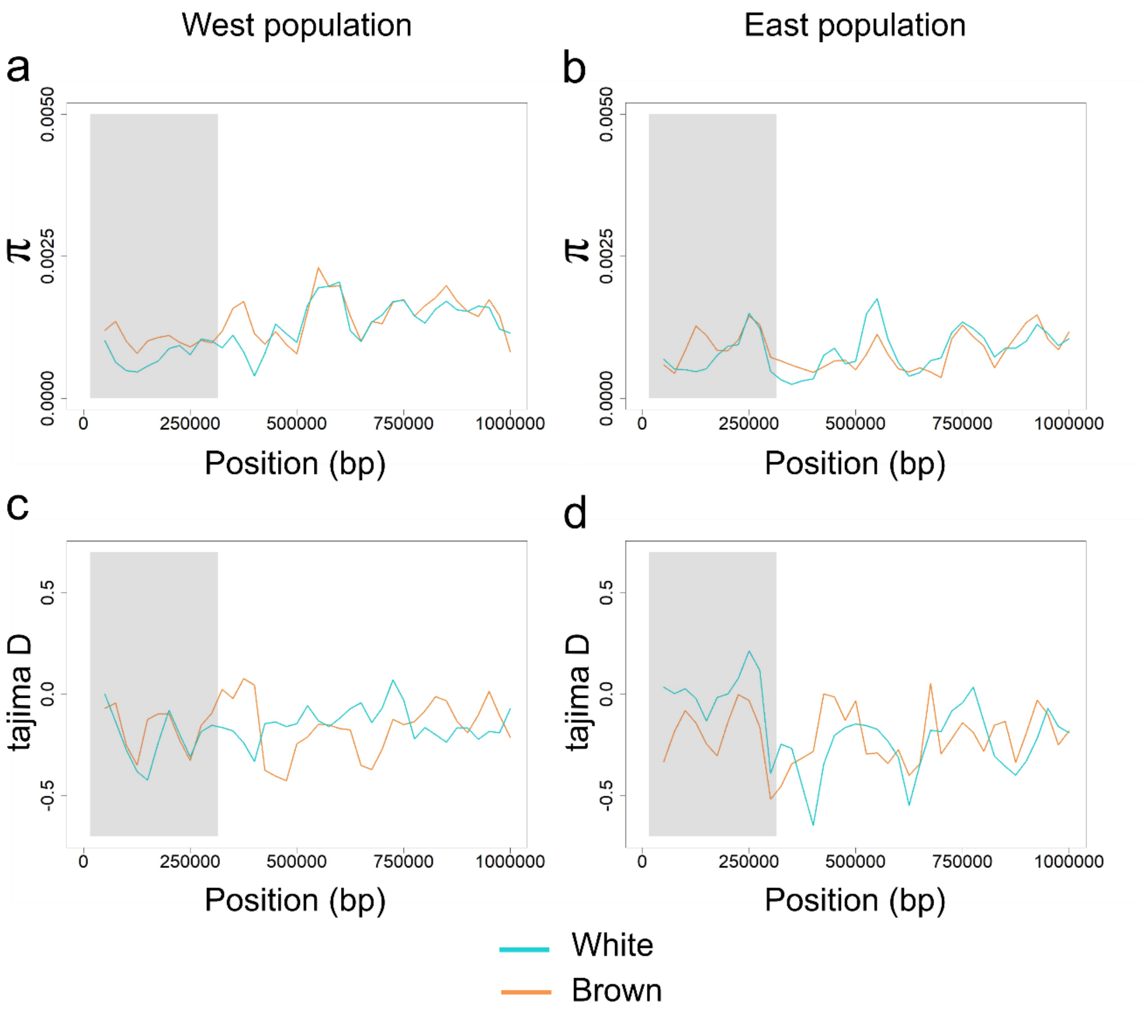


**Figure S6. Nucleotide diversity and Tajima's D along the first 1 Mb of chromosome 7.** Nucleotide diversity was inferred in overlapping windows of 50 kb (sliding 25 kb) for each colour morph from West (a) and East (b) populations. Tajima D estimations inferred in overlapping windows of 50 kb (sliding 25 kb) for each colour morph from West (c) and East (d) populations. The grey shaded area shows the winter polymorphism association region.

a

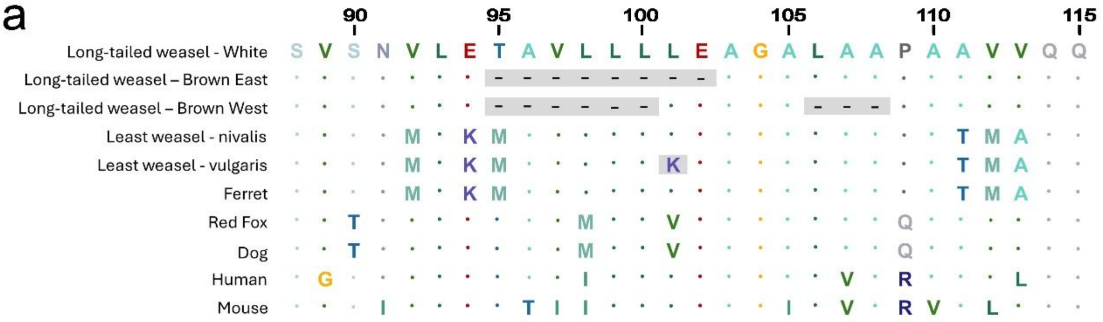


b

| Substitution Type | Region | Amino acid position change | Effect | Confidence Score |
|---|---|---|---|---|
| AA_DELETION | CDS | 94-102 | damaging | 0.894 |
| AA_DELETION | CDS | 94-100 | damaging | 0.894 |
| AA_DELETION | CDS | 106-109 | damaging | 0.894 |

c

| Chromosome | Position | Reference | Alt | RawScore | PHRED |
|---|---|---|---|---|---|
| 16 | 89985945 | GGAGACGGCCGTCATCCTC | G | 1.579225 | 16.73 |
| 16 | 89985945 | GGAGACGGCCGTCATCCTCCTGCTG | G | 1.553943 | 16.58 |

**Figure S7. Candidate mutations, functional impact and population structure.**
(a) Multispecies alignment of MC1R sequences around the structural variants identified in the protein. Shaded grey segments highlight candidate variants to underlie winter colour polymorphism in winter brown long-tailed and least weasels. (b-c) *In silico* tests of the functional impact of long-tailed weasel deletions using (b) SIFT, showing a confidence score of 0.894 for damaging impact in gene function, and (c) CADD, showing a scaled score (PHRED) of >16 for the largest structural variants detected that place them in the top 10% of the mutations with deleterious effect for the human genome.
